# Ceramide transfer protein regulates G-protein coupled phospholipase signalling in *Drosophila* photoreceptors

**DOI:** 10.1101/2025.08.05.668821

**Authors:** Shirish Mishra, Ali Khan, Vaisaly R Nath, Tejaswini Manoj, BG Thejaswini, Amruta Naik, Ujjaini Dasgupta, Padinjat Raghu

**Author notes:** Corresponding authors: Padinjat Raghu, Shirish Mishra.

## Abstract

The non-vesicular transfer of lipids between organelles at membrane contact sites (MCS) has been proposed as a key principle in the regulation of cell physiology. While several proteins with lipid transfer activity have been identified and localized to MCS, their functional significance for supporting physiology is poorly understood. Ceramide transfer protein (CERT) is one such molecule that can transfer ceramide between membranes *in vitro*. However, evidence for the mechanism and *in vivo* significance of CERT function is limited. In this study, we have analyzed the function of the only gene (*dcert*) encoding CERT in *Drosophila*. We find that loss of function alleles of *dcert* (*dcert^1^*), show elevated levels of short chain ceramide species along with a reduction in the levels of its metabolite phosphatidyl ethanolamine ceramide. Physiological analysis of *dcert^1^* mutant alleles showed reduced electrical responses in the eye to light stimulation although photoreceptors did not undergo retinal degeneration, and this phenotype could be rescued by reconstitution of *dcert^1^* with the wild type gene. The altered light response in dcert*^1^* was associated with a reduction in the rate of phosphatidylinositol 4,5 bisphosphate (PIP_2_) resynthesis following light induced phospholipase C (PLC) stimulation. The reduced electrical response of *dcert^1^* could be suppressed by reducing ceramide synthesis at the ER. Taken together, our findings suggest that ceramide synthesized at the ER and transferred to the Golgi by CERT regulates G-protein coupled phospholipase C signaling *in vivo*.

## Introduction

Lipids are key biomolecular components of cells and serve multiple functions including structural and signalling roles. In eukaryotic cells which contain membrane bound organelles, lipid synthesis occurs primarily in the smooth endoplasmic reticulum (sER) from which they are transported to organelle membranes where they or their metabolites serve specific functions. Lipid transport and exchange between eukaryotic organelle membranes can occur through a variety of mechanisms; these include both vesicular transport as well as the activity of proteins, referred to as lipid transfer proteins (LTP) that are able to transfer lipids from one organelle membrane to another. A subset of these LTPs are localized to membrane contact sites (MCS) and their function at these MCS(Cockcroft and Raghu, 2018) underscores the importance of lipid transfer between organelle membranes without membrane fusion.

Ceramide transfer protein (CERT), also known as Goodpasture Antigen Binding Protein, is one such LTP that is localized to the MCS between the endoplasmic reticulum (ER) and the Golgi apparatus(Hanada et al., 2003; Yamaji and Hanada, 2015). CERT transfers ceramide from the sER, the primary site of ceramide synthesis to the Golgi apparatus where it is used for sphingomyelin synthesis. Thus, CERT function is critical for regulated sphingolipid synthesis. This finding is underscored by the report that deletion of CERT in mice results in lethality during embryonic development(Wang et al., 2009), depletion in flies results in reduced lifespan (Rao et al., 2007) and mutations in critical residues of CERT have been associated with intellectual disability in human patients(Gehin et al., 2023; de Ligt et al., 2012; Hamdan et al., 2014; Deciphering Developmental Disorders Study, 2015, 2017; Murakami et al., 2020). At a cellular level, CERT has been implicated in mitochondrial function(Wang et al., 2009; Rao et al., 2007) and plasma membrane (PM) organization(Kunduri et al., 2018). In principle, loss of CERT activity can result in multiple biochemical changes including the accumulation of ceramides in the ER as well as reduction of sphingomyelin and other sphingolipid metabolites at the PM or other membranes following transport of ceramides out of the ER by CERT. Consistent with this, numerous studies have noted the importance of the ceramide/sphingomyelin ratio for the behavior of cancer cells and therefore of clinical outcomes in cancer(Pal et al., 2022; Piazzesi et al., 2021; Li et al., 2022). Therefore, the biochemical mechanism underlying the phenotypic outcomes of CERT loss of function *in vivo* is less clear.

*Drosophila* photoreceptors are large polarized cells whose apical domain is expanded and specialized to detect and transduce photons of light using G-protein coupled phospholipase C signalling(Raghu et al., 2012). Previous studies using this model have highlighted the role of sphingolipid metabolism in photoreceptors homeostasis. Loss of function in ceramide kinase, a key enzyme mediating the signalling function of ceramide result in a reduced light response and photoreceptor degeneration(Dasgupta et al., 2009). Further, overexpression of ceramidase in *Drosophila* photoreceptors has been reported to modulate retinal degeneration in multiple mutants including *norpA^P24^* (phospholipase C), *arr2^3^*(arrestin) and *ninaE^117^* (Rhodopsin 1)(Acharya et al., 2003, 2008, 2004). However, a strong hypomorph in *Drosophila* CERT (*dcert)* was not reported to show neuronal degeneration(Rao et al., 2007) and its impact on photoreceptor structure and function is not known although in a recent study we noted that RNAi depletion of *dcert* can modulate the retinal degeneration phenotype of *rdgB* and also noted an altered electrical response to light(Mishra et al., 2024). Here we report that the strong hypomorph *dcert^1^*shows a reduced electrical response to light at eclosion that further declines with age without concomitant retinal degeneration. This defective light response was associated with altered turnover of phosphatidylinositol 4 phosphate (PI4P) and phosphatidylinositol 4,5 bisphosphate (PIP_2_) at the PM, underpinned by defects in photoreceptor ER-PM MCS. Biochemical analysis of *dcert^1^* revealed a reduction in multiple species of photoreceptor phosphoethanolamine-ceramides (PE-Cer) associated with an elevation in the levels of some species of ceramides. Our genetic and biochemical analysis suggests that altered PM function, dependent on PE-Cer, plays a key role in the altered light response in *dcert^1^*. Overall, our studies reveal the importance of CERT dependent, PM PE-Cer in the regulation of ER-PM MCS and G-protein coupled PLC signalling *in vivo*.

## Results

### A protein null mutant (*dcert^1^*) shows reduced ERG amplitude but no retinal degeneration

An earlier study had noted that depletion of *dcert* by RNAi resulted in a modest reduction in the electrical response to light measured through electroretinogram (ERG)(Mishra et al., 2024). To confirm this result, we studied *dcert^1^*, a previously described protein null allele in this gene(Rao et al., 2007). Western blot analysis of *dcert^1^* confirmed no detectable full-length DCERT protein (**Fig 1A**). ERG recordings revealed that *dcert^1^* photoreceptors showed a reduced electrical response to light (**Fig 1 B, C**). This reduced ERG amplitude could be rescued by reconstituting *dcert^1^* with a bacterial artificial chromosome(Venken et al., 2006) containing the *dcert* gene (**Fig 1B, C**) demonstrating that the reduced ERG amplitude in *dcert^1^* results from the loss of *dcert*. To corroborate our findings, we also placed the *dcert^1^* allele in trans with a deficiency *Df(732)* that deletes *dcert*. *dcert^1^*/*Df(732)* flies showed a reduced ERG amplitude (**Fig 1 D, E)** like that seen for *dcert^1^* homozygous flies. Following eclosion we reared *dcert^1^* for ten days and found that photoreceptors did not undergo retinal degeneration (**Fig 1 F)**. Further, *dcert^1^* reared in constant white light illumination (2800 lux) for ten days also did not undergo retinal degeneration (data not shown). However, interestingly and in contrast to controls, the ERG amplitude of *dcert^1^* flies showed a progressive reduction in amplitude as a function of age (Fig 1G). These findings suggest that *dcert* function is required to support a normal electrical response to light.

**Figure 1:**
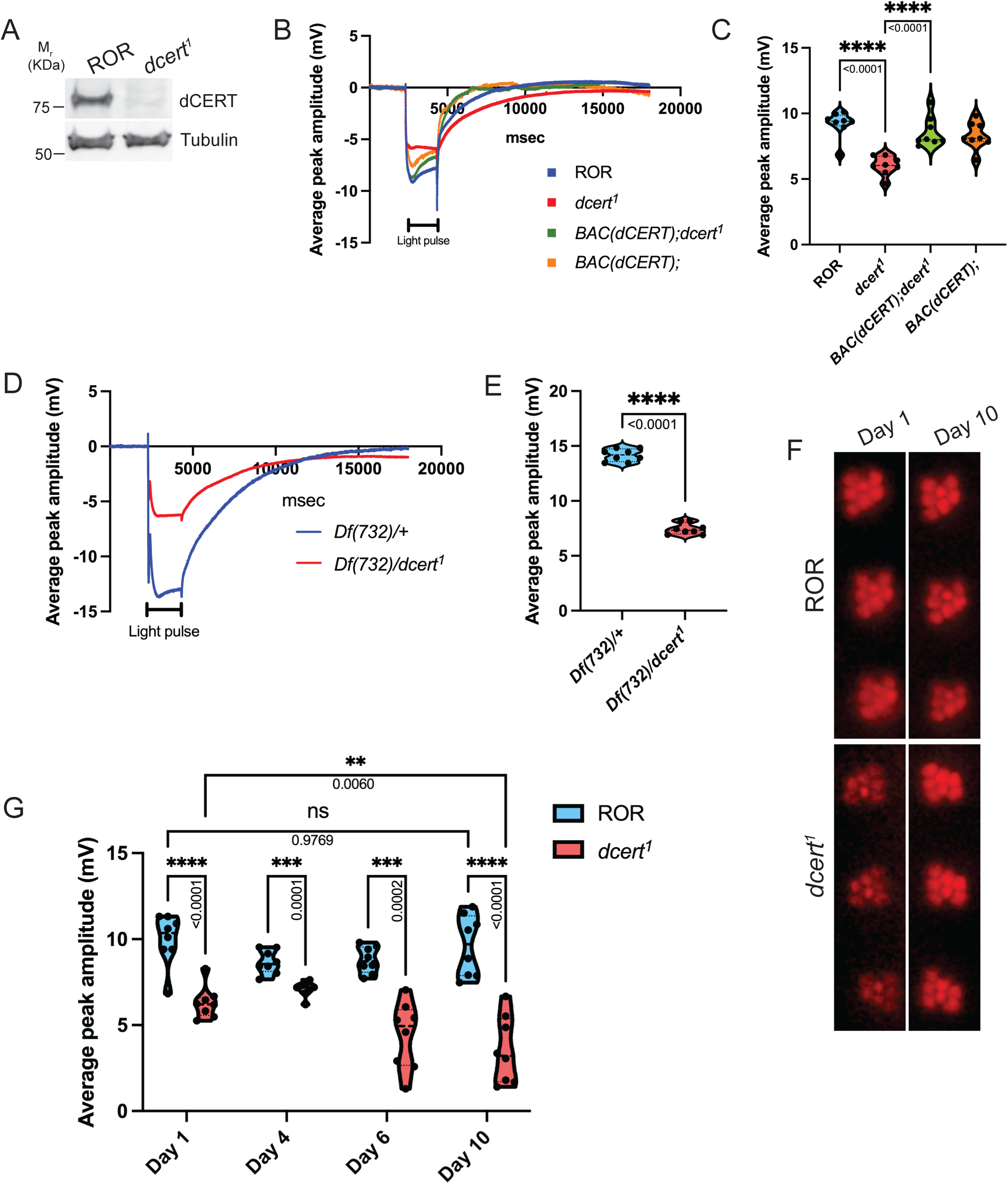
dCERT regulates light response in *Drosophila* photoreceptors: (A) Western blot analysis from fly head extract of indicated genotypes is shown. The blot was probed with anti-dCERT antibody. Tubulin was used as a loading control. (B) Representative ERG traces from dark reared 0 to 1 day old flies indicating the response of control (ROR), *dcert^1^*, *BAC(dCERT); dcert^1^* and BAC(dCERT) photoreceptors to 2s flash of green light. BAC(dCERT) represents BAC clone containing *dCERT* gene. The duration of the stimulating light is shown. X axis shows the duration of recording (msec) while y axis shows the amplitude (mV). (C) Quantification of ERG amplitude of indicated genotypes. Each data point depicts a single fly tested (n=10 biological replicates). Violin plots with the mean ± SEM are shown. Statistical tests: 2 way ANNOVA grouped analysis. (D) Representative ERG traces from dark reared 0 to 1 day old flies indicating the response of heterozygous deficiency line 732/+ (*Df732/+*) and *Df732/dcert^1^*. The duration of the stimulating light is shown. X axis shows the duration of recording (msec) while y axis shows the amplitude (mV). (E) Quantification of ERG amplitude of indicated genotypes. Each data point depicts a single fly tested (n=10 biological replicates). Violin plots with the mean ± SEM are shown. Statistical tests: Student’s unpaired t test. (F) Representative optical neutralization (ON) images depicting rhabdomere integrity of indicated genotypes when reared in dark. Images were taken on day 1 and day 10. (G) Quantification of ERG amplitude of indicated genotypes. Each data point depicts a single fly tested (n=10 biological replicates). This result shows gradual decrease in ERG amplitude in *dcert^1^* with age. Violin plots with the mean ± SEM are shown. Statistical tests: 2 way ANNOVA grouped analysis.

### *dcert* is required for phosphoinositide turnover during PLC signalling

In *Drosophila* photoreceptors, sensory transduction is mediated by the G-protein coupled receptor (GPCR) Rhodopsin 1 (Rh1) that activates a G_q_ coupled PLC activity, leading to opening of TRP and TRPL channels(Raghu et al., 2012). To understand the mechanism underlying the reduced ERG amplitude in *dcert^1^*, we performed Western blot analysis to check the levels of the major phototransduction proteins. This analysis revealed no reduction in the levels of Rh1, that detects light in photoreceptors (**Fig 2A**). The levels of other major phototransduction proteins such as NORPA, INAD, TRP and Arrestin 2 were not altered in *dcert^1^* (**Fig 2A**). An alternate possibility for the reduced ERG response is that the levels of the lipid substrate of NORPA, PIP_2_ is reduced in *dcert^1^*. We measured the levels of PIP_2_ at the photoreceptor PM using the fluorescent PIP_2_ reporter PH-PLC8::GFP (Várnai and Balla, 1998; Chakrabarti et al., 2015). Under resting conditions, we found a modest reduction in the levels of the probe at the apical PM of *dcert^1^* implying a reduction in PIP_2_ levels (**Fig 2B**). We also monitored the turnover of the probe at the apical PM following stimulation with a bright light that depletes all the PIP_2_ at this location. Under these conditions, we found that the rate of recovery of PIP_2_ levels was slower in *dcert^1^* compared to controls (**Fig 2C**). Low PIP_2_ can be attributed to reduced protein levels of dPIP5K, enzyme that synthesizes PIP_2_ at the PM during phototransduction(Chakrabarti et al., 2015; Kumari et al., 2022). We found that the levels of dPIP5K were unchanged in *dcert^1^* (**Fig 2D**). Similar analysis using a probe that detects PM pools of PI4P(Hammond et al., 2014; Balakrishnan et al., 2018) revealed no change in the levels of PI4P under resting conditions (**Fig 2E**); however the rate at which PI4P was resynthesized following depletion by a bright flash of light was slower in *dcert^1^* compared to controls (**Fig 2F**) even though mRNA levels of PI4P synthesizing enzyme dPI4KIIIα(Balakrishnan et al., 2018) was higher in *dcert^1^* (**Fig 2G**).

**Figure 2:**
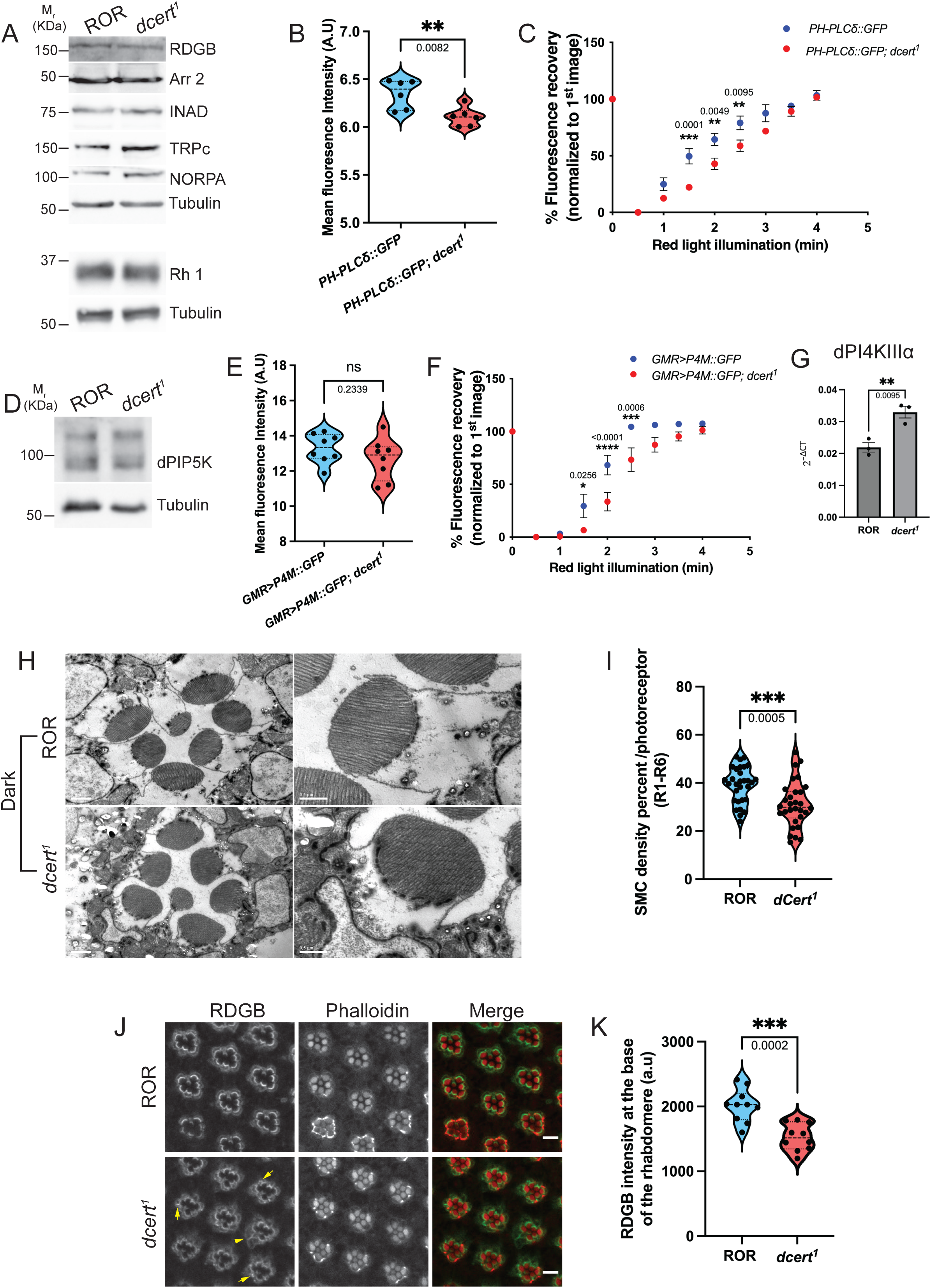
Loss of dCERT leads to reduced PI(4,5)P_2_ recycling and MCS defects: (A) Western blot from head extract of indicated genotypes was probed with antibodies of major phototransduction proteins. Tubulin is used as the loading control. (B) Quantification of the mean fluorescence intensity of the PI(4,5)P_2_ probe PH-PLC8-GFP from the deep pseudopupil formed by one-day-old flies of the indicated genotypes. (n=6 biological replicates). (C) Recovery kinetics of the PI(4,5)P_2_ probe PH-PLC8-GFP from fluorescence deep pseudopupil over time. Average fluorescence intensity of each pseudopupil image is expressed as a percentage of the intensity of first image acquired. Each data point represents mean ± SEM. (n=6 biological replicates) Statistical tests: 2 way ANNOVA grouped analysis. (D) Western blot analysis from fly head extract of indicated genotypes is shown. The blot was probed with anti-dPIP5K antibody. Tubulin was used as a loading control. (E) Quantification of the mean fluorescence intensity of the PI4P probe P4M-GFP from the deep pseudopupil formed by one-day-old flies of the indicated genotypes. (n=8 biological replicates). (F) Recovery kinetics of the PI4P probe P4M-GFP from fluorescence deep pseudopupil over time. Average fluorescence intensity of each pseudopupil image is expressed as a percentage of the intensity of first image acquired. Each data point represents mean ± SEM. (n=8 biological replicates) Statistical tests: 2 way ANNOVA grouped analysis. (G) Quantitative real time PCR showing mRNA transcript levels of dPI4KIIIα in adult fly retinae of indicated genotypes (n=3 biological replicates with 80 retinae/biological replicate). Transcript level expression was normalised to the loading control, ribosomal protein 49 (RP49). Statistical tests: Student’s unpaired t test. (H) TEM images of single ommatidia (scale bar: 1μm) and magnified photoreceptors (scale bar: 0.5μm) from ROR and *dcert^1^*of 1-day old dark reared flies. (I) Quantification of the SMC density per photoreceptor in indicated genotypes. n =30 photoreceptors from 3 separate flies for R1-R6. Statistical tests: Student’s unpaired t test. (J) Confocal images showing localization of RDGB in ROR and *dcert^1^* photoreceptors of 1-day old dark reared flies. RDGB puncta are depicted here by arrows and solid triangle show presence of RDGB in the cell body. Rhabdomeres are stained with phalloidin that binds to F-actin. Scale bar: 5μm. (K) Quantification of RDGB intensity at the base of the rhabdomere in ROR and *dcert^1^* photoreceptors. RDGB intensity using line plot was analysed in 60 photoreceptors from two independent retinae in each genotype. n=10 represents 10 ommatidia where each ommatidia has R1-R6 light sensitive photoreceptors. Statistical tests: Student’s unpaired t test.

### ER-PM MCS defects in *dcert^1^* photoreceptors

In *Drosophila* photoreceptors PM-sub microvillar cisternae (SMC) MCS are required to support PIP_2_ resynthesis for sustained signaling (Yadav et al., 2015)[reviewed in (Yadav et al., 2016)]. To determine if the reduced rate of PI4P and PIP_2_ synthesis in *dcert^1^* was due to defects in ER-PM MCS, we visualized and quantified these by transmission electron microscopy. This analysis revealed that at eclosion, the density of ER-PM MCS in *dcert^1^* was lower than in controls (**Fig 2 H, I)** and somewhat exacerbated by light exposure (**Sup figure 2A, B).** Reduced ER-PM MCS density could lead to improper localization of proteins normally restricted to this location; RDGB, the multidomain phosphatidylinositol transfer protein is one such protein(Yadav et al., 2015, 2016; Vihtelic et al., 1993). We found the levels of RDGB protein were no different between controls and *dcert^1^* (**Fig 2A**). However, immunolocalization studies revealed that while RDGB is strictly localized to the ER-PM MCS in wild type cells, in *dcert^1^*, a significant fraction of RDGB protein is mislocalised away from the ER-PM MCS and formed puncta (**Fig 2J, K)**.

### Altered ceramide and phosphoethanolamine ceramide levels in *dcert^1^*

To understand the biochemical basis of the altered phototransduction phenotype in *dcert^1^*, we measured the levels of sphingolipids in *Drosophila* head extracts using liquid chromatography (LC) coupled with tandem mass spectrometry (MS). We found that overall, total ceramide levels did not show any significant change in the *dcert^1^*mutant in comparison to wild type (**Fig 3A**). However, an inspection of the levels of individual ceramide molecular species revealed that levels of ceramide species with shorter chain length in d-14 and d-16 series such as d14:c14, d14:c16, d16:c16 ceramide were significantly higher while the levels of d14:c18 and d16:c18 were lower in *dcert^1^* than wild type (**Fig 3B**). The most abundant ceramide species c-20 and c-22 in d-14 ceramide series did not show any difference between mutant and wild type (**Fig 3B**).

**Figure 3:**
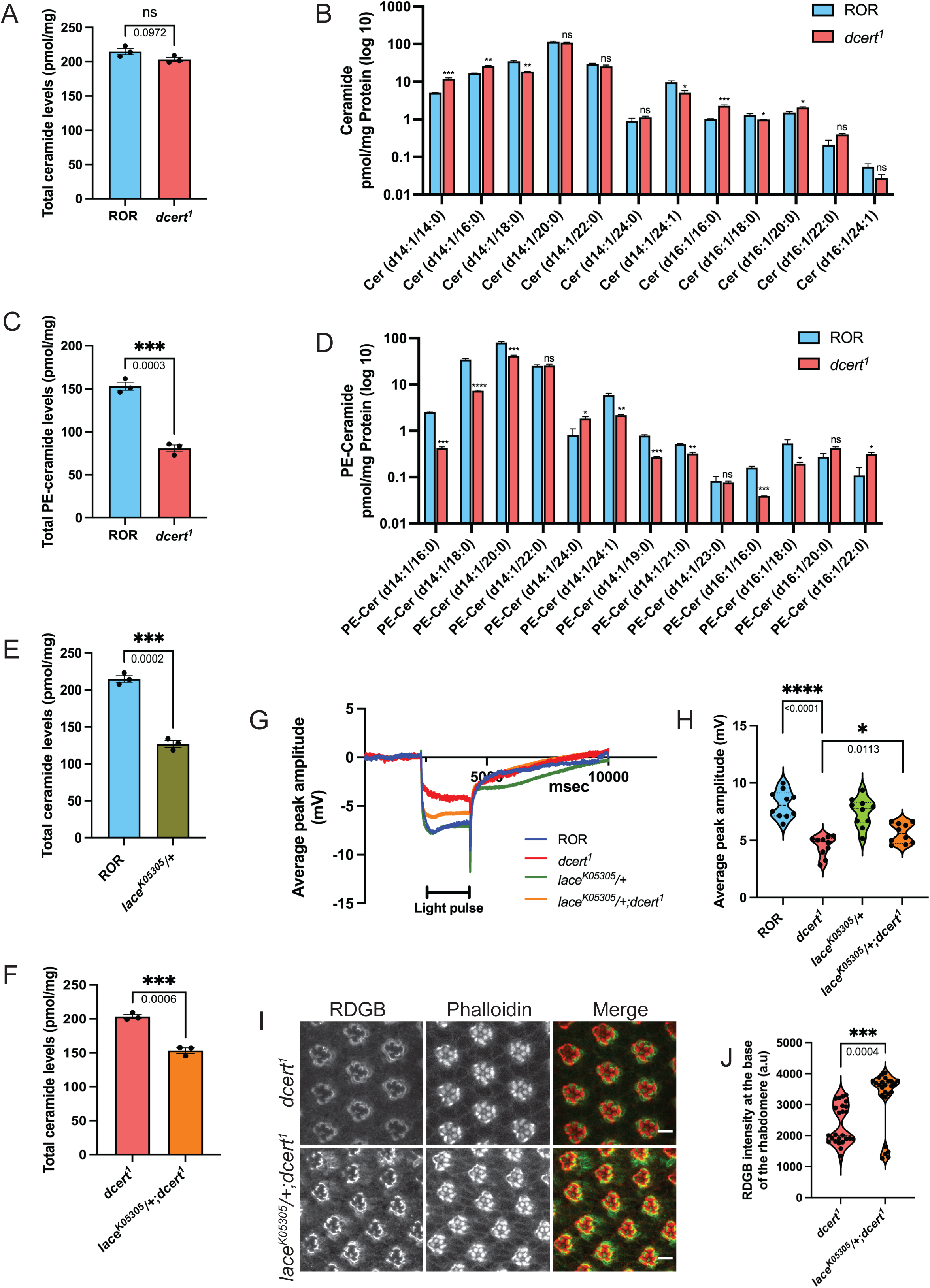
Reduced ceramide formation at the ER partially rescues ERG phenotype in *dcert^1^*: (A) Total ceramide levels in adult fly heads in ROR and *dcert^1^*. Y axis shows level of ceramide as pmol/mg of the total protein present in the sample. Statistical tests: Student’s unpaired t test. n=3. (B) Comparison of individual ceramide species in adult fly heads in ROR and *dcert^1^*. Y axis shows level of ceramide as pmol/mg of the total protein present in the sample in the log_10_ scale. Statistical tests: Student’s unpaired t test for each species. (C) Total phosphoethanolamine-ceramide (PE-ceramide) levels in adult fly heads in ROR and *dcert^1^*. Y axis shows level of PE-ceramide as pmol/mg of the total protein present in the sample. Statistical tests: Student’s unpaired t test. n=3. (D) Comparison of individual PE-ceramide species in adult fly heads in ROR and *dcert^1^*. Y axis shows level of PE-ceramide as pmol/mg of the total protein present in the sample in the log_10_ scale. Statistical tests: Student’s unpaired t test for each species. (E) Total ceramide levels in adult fly retinae in ROR and lace*^K05305^/+*. Y axis shows level of ceramide as pmol/mg of the total protein present in the sample. Statistical tests: Student’s unpaired t test. n=3. (F) Total ceramide levels in adult fly retinae in *dcert^1^* and lace*^K05305^/ dcert^1^*. Y axis shows level of ceramide as pmol/mg of the total protein present in the sample. Statistical tests: Student’s unpaired t test. n=3. (G) Representative ERG traces from dark reared 0 to 1 day old flies indicating the response of control (ROR), *dcert^1^*, lace*^K05305^/+* and lace*^K05305^/ dcert^1^* photoreceptors to 2s flash of green light. The duration of the stimulating light is shown. X axis shows the duration of recording (msec) while y axis shows the amplitude (mV). (H) Quantification of ERG amplitude of indicated genotypes. Each data point depicts a single fly tested (n=10 biological replicates). Violin plots with the mean ± SEM are shown. Statistical tests: Student’s unpaired t test. (I) Confocal images showing localization of RDGB in *dcert^1^* and lace*^K05305^/ dcert^1^* photoreceptors of 1-day old dark reared flies. Rhabdomeres are stained with phalloidin that binds to F-actin. Scale bar: 5μm. (J) Quantification of RDGB intensity at the base of the rhabdomere in *dcert^1^* and lace*^K05305^/ dcert^1^*photoreceptors. RDGB intensity using line plot was analysed in 150 photoreceptors from five independent retinae in each genotype. n=25 represents 25 ommatidia where each ommatidia has R1-R6 light sensitive photoreceptors. Statistical tests: Student’s unpaired t test.

In *Drosophila*, phosphoethanolamine-ceramides (PE-Cer) are formed in Golgi by the action of the enzyme ceramide phosphoethanolamine synthase(Acharya and Acharya, 2005) encoded by the genes *CPES*(Kunduri et al., 2018) and sphingomyelin Synthase related (SMSr)(Vacaru et al., 2009). While CPES generates the majority of PE-Cer through transfer of the CDP-ethanolamine as headgroup donor to a ceramide molecule, SMSr forms trace amount of PE-ceramide. The substrate for these enzymes, ceramide is transferred from the site of synthesis, the ER to the Golgi apparatus by the lipid transfer activity of CERT. This predicts that in mutants lacking CERT activity, PE-Cer levels might be reduced. We measured PE-Cer levels and found it to be reduced by ca. 50 % in *dcert^1^*(**Fig 3C**). When individual PE-Cer species were quantified both short and long chain species in d-14 (except c-24) and d-16 series (except c-22) were found significantly lower in the mutant than in the wild type (**Fig 3D**). Overall, depletion of *dcert* results in an elevation of some Cer species along with a concomitant reduction of most species of PE-Cer.

### Elevated ceramide levels contribute to the reduced light response in *dcert^1^*

When CERT function is abrogated in *dcert^1^,* ceramide levels are expected to rise at the sER, the site of its synthesis. To test if buildup of ceramide in the ER is responsible for the reduced light response in *dcert^1^,* we tested the effect of reducing ER ceramide levels in *dcert^1^*. Ceramide synthesis is initiated by the activity of the rate limiting enzyme, serine palmitoyl transferase (SPT), encoded in flies by *lace*(Adachi-Yamada et al., 1999). *lace* allele *l(2)k05305* contains a P-element insertion and is homozygous lethal. We checked levels of ceramide in *lace^k05305^ /+* heterozygous mutant heads using LC-MS and found ca. 50% reduction in its levels (**Fig 3E, Sup.** Fig 2A). We then generated a *lace^k05305^*/+; *dcert^1^* double mutant; LC-MS analysis of head extracts from these flies revealed that total ceramide levels in the double mutant were significantly reduced compared to *dcert^1^* (**Fig 3F, Sup.** Fig 2B).

Therefore, reduction of SPT activity in *dcert^1^* can lower elevated ceramide levels. To test the effect of reducing ceramide levels in *dcert^1^*, we measured the ERG amplitude of *lace^k05305^* /+; *dcert^1^*. This analysis revealed that the ERG amplitude of *lace^k05305^* /+; *dcert^1^* was modestly elevated compared to *dcert^1^*alone (**Fig 3 G, H)**; *lace^k05305^* /+ flies themselves had no reduction in ERG amplitude and flies where lace mRNA levels were downregulated in the eye using RNAi did not show reduction in the ERG amplitude **(Sup.** Fig 2C**, 2D)**. Further, we found that the levels of RDGB protein at the ER-SMC contact sites, reduced in *dcert^1^*, were elevated in *lace^k05305^* /+; *dcert^1^* (**Fig 3 I, J)**. These findings suggest that elevation of ceramide in the ER contributes partially to the reduced ERG amplitude and reduced levels of RDGB at the SMC in *dcert^1^*.

### Reduced PE-Cer levels result in a reduced electrical response to light

Since reducing ceramide levels in the ER only partially suppressed ERG phenotype of *dcert^1^*, we tested the possibility that the reduced levels of PE-Cer may be responsible for the defective light response. CPES activity is encoded by two genes *CPES* and *SMSr* (**Fig 4A**). We selectively downregulated CPES and SMSR selectively in the eye using RNAi. Using LC-MS, levels of ceramide in the *GMR>CPES^i^*were increased while that of PE-ceramide were reduced in fly retinae (**Fig 4D, 4E, Sup.** Fig 3B**, 3C)**. ERG analysis revealed that knockdown of CPES mRNA and not of SMSr mRNA showed a reduced ERG amplitude (**Fig 4B, 4C, Sup.** Fig 3A) compared to controls. This finding phenocopies the reduced ERG amplitude of *dcert^1^*. Further, Western blot analysis revealed that like *dcert^1^*, the levels of RDGB did not change in *GMR>CPES^i^* (**Fig 4F**) although its localization at the base of the rhabdomere is highly reduced and protein is either present in the cell body as puncta (**Fig 4G, 4H)**. These findings suggest an important contribution for reduced PE-Cer levels in the eye phenotypes of *dcert^1^*.

**Figure 4:**
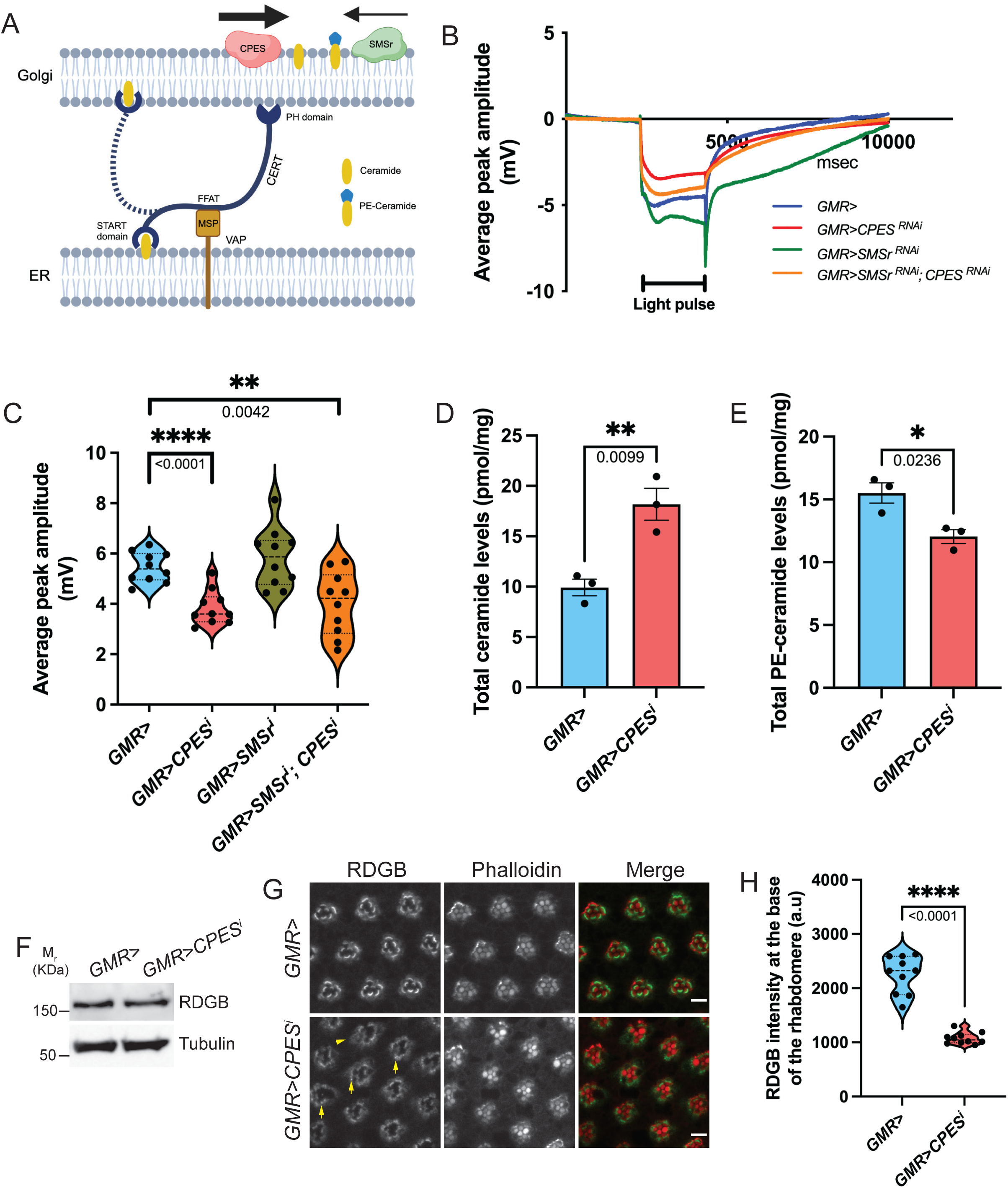
Flies with reduced CPES activity mimic *dcert^1^* phenotypes: (A) Cartoon depicting dCERT dependent flow of ceramide from ER to the Golgi where two enzymes Ceramide phosphoethanolamine synthase (CPES) and Sphingomyelin synthase related (SMSr) catalyse the formation of PE-Ceramide at the Golgi. Figure also depicts that majority of the PE-ceramide at Golgi is formed by CPES enzyme (thick arrow). (B) Representative ERG traces from dark reared 0 to 1 day old flies indicating the response from indicated genotypes to 2s flash of green light. CPES and SMSr knockdown was achieved using strong eye promoter, GMR. The duration of the stimulating light is shown. X axis shows the duration of recording (msec) while y axis shows the amplitude (mV). (C) Quantification of ERG amplitude of indicated genotypes. Each data point depicts a single fly tested (n=10 biological replicates). Violin plots with the mean ± SEM are shown. Statistical tests: Student’s unpaired t test. (D) Total ceramide levels in adult fly retinae in GMR> and *GMR>CPES^i^*. Y axis shows level of ceramide as pmol/mg of the total protein present in the sample. Statistical tests: Student’s unpaired t test. n=3. (E)Total PE-ceramide levels in adult fly retinae in GMR> and *GMR>CPES^i^*. Y axis shows level of ceramide as pmol/mg of the total protein present in the sample. Statistical tests: Student’s unpaired t test. n=3. (F) Western blot analysis from fly head extract of indicated genotypes is shown. The blot was probed with anti-RDGB antibody. Tubulin was used as a loading control. (G) Confocal images showing localization of RDGB in GMR> and *GMR>CPES^i^*photoreceptors of 1-day old dark reared flies. RDGB puncta are depicted here by arrows and solid triangle show presence of RDGB in the cell body. Rhabdomeres are stained with phalloidin that binds to F-actin. Scale bar: 5μm. (H) Quantification of RDGB intensity at the base of the rhabdomere in GMR> and *GMR>CPES^i^* photoreceptors. RDGB intensity using line plot was analysed in 60 photoreceptors from two independent retinae in each genotype. n=10 represents 10 ommatidia where each ommatidia has R1-R6 light sensitive photoreceptors. Statistical tests: Student’s unpaired t test.

### Altered PM organization in *dcert^1^*

To test if PM protein composition is altered in *dcert^1^*, we measure the levels of several key proteins involved in *Drosophila* phototransduction that are located at the apical PM or closely associated with it. To test if the localization of phototransduction proteins was normal we compared the localization of Rh1, TRP and INAD, a key molecule involved in the organization of the transduction complex at the PM(Chevesich et al., 1997; Tsunoda et al., 1997). This analysis revealed that there was no difference in the localization of TRP (**Fig 5A**), INAD (**Fig 5B**) and Rh1 (**Fig 5C**).

**Figure 5:**
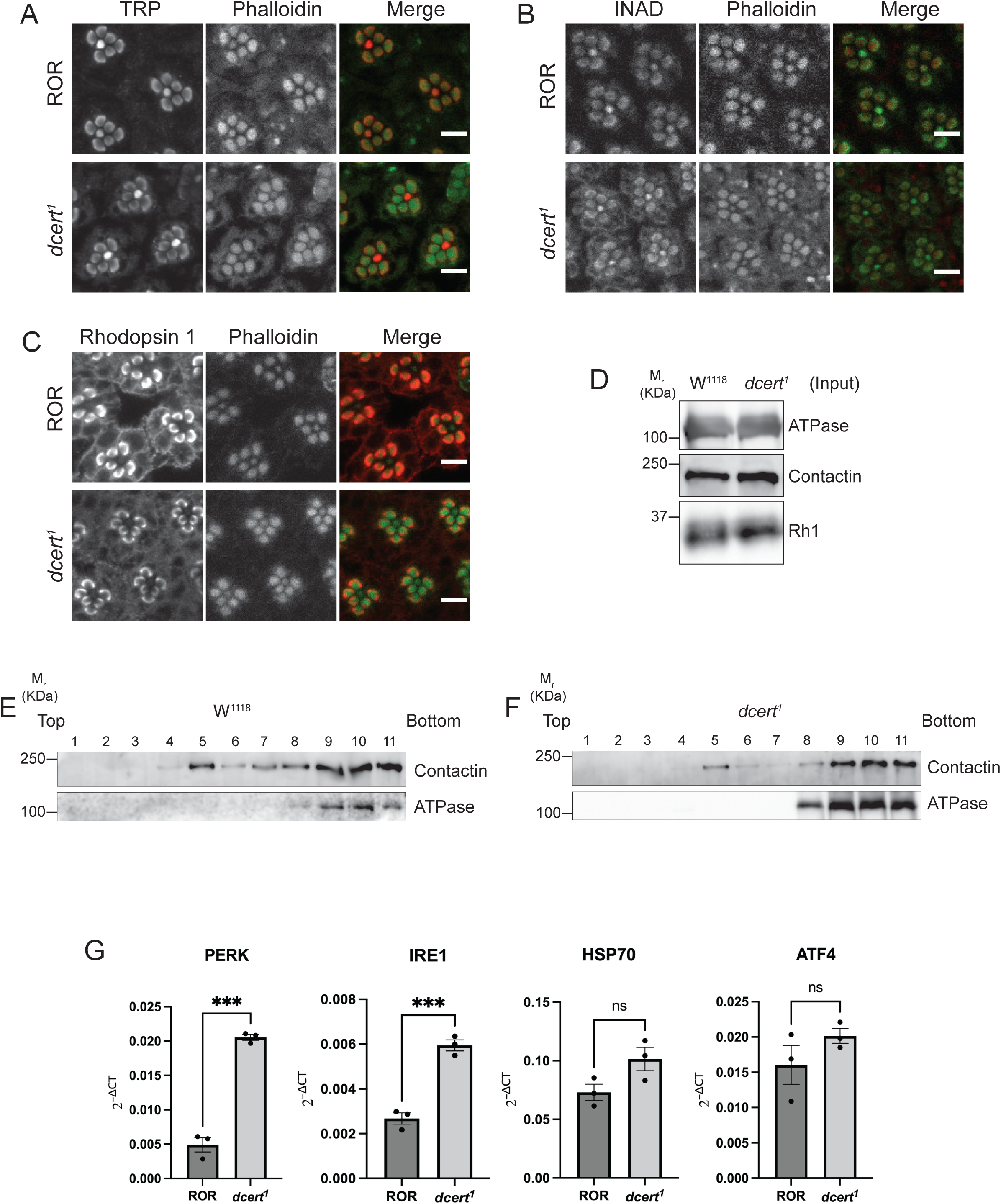
*dcert^1^*has defects in plasma membrane lipid ordering and high UPR: (A) Confocal images showing localization of TRP in ROR and *dcert^1^* photoreceptors of 1-day old dark reared flies. Rhabdomeres are stained with phalloidin that binds to F-actin. Scale bar: 5μm. (B) Confocal images showing localization of INAD in ROR and *dcert^1^* photoreceptors of 1-day old dark reared flies. Rhabdomeres are stained with phalloidin that binds to F-actin. Scale bar: 5μm. (C) Confocal images showing localization of rhodopsin 1 (Rh1) in ROR and *dcert^1^* photoreceptors of 1-day old dark reared flies. Rhabdomeres are stained with phalloidin that binds to F-actin. Scale bar: 5μm. (D) Detergent resistant membranes (DRMs) were isolated from fly heads. Input was quantified by Bradford and equal amount of protein was loaded. Blot was analysed for ATPase, Contactin and Rh1. (E) Equal amount of protein was loaded in each fraction from top to bottom of the sucrose gradient and analysed by western blotting in W^1118^. The blot was probed with contactin and ATPase. (F) Equal amount of protein was loaded in each fraction from top to bottom of the sucrose gradient and analysed by western blotting in *dcert^1^*. The blot was probed with contactin and ATPase. (G) Quantitative real time PCR showing mRNA transcript levels of PERK, IRE1, HSP70 and ATF4 in adult fly retinae of indicated genotypes (n=3 biological replicates with 80 retinae/biological replicate). Transcript level expression was normalised to the loading control, ribosomal protein 49 (RP49). Statistical tests: Student’s unpaired t test.

PM sphingolipids have been proposed to play a key role in the organization of detergent resistant domains (DRMs) at the PM of cells(Rietveld et al., 1999) (Simons and Sampaio, 2011). To test if the loss of *dcert* function affects the organization of these DRMs, we fractionated membrane lysates from wild type and *dcert^1^* heads using a sucrose density gradient(Kunduri et al., 2018). We assayed fractions for the presence of DRMs using antibodies for contactin, a GPI anchored septate junction protein(Faivre-Sarrailh et al., 2004) known to be localized to these domains. Na-K ATPase, a non-DRM protein; Contactin and Rh1 were used as controls to ensure that the starting lysates had equal amounts of protein (**Fig 5E**). Using this approach, we found that while contactin was enriched in DRM fractions 5-8 in control flies (**Fig 5F**), presence of contactin is highly reduced in these fractions in *dcert^1^* (**Fig 5G**). These findings suggest that the levels of DRMs at the PM are reduced in *dcert^1^*mutants.

### Unfolded Protein response upregulated in *dcert^1^*

Previous studies have shown that lipid accumulation in the ER can lead to ER stress and upregulation of unfolded protein response (UPR)(Celik et al., 2023). To test whether altered ceramide metabolism leads to ER stress we measured the levels of four UPR genes in *dcert^1^*. These include inositol requiring enzyme 1 (IRE1), double stranded RNA activated protein kinase (PKR)-like ER kinase (PERK), heat shock protein 70 (HSP70) and transcription factor ATF4. IRE1 and PERK are highly conserved ER stress sensors and relay downstream signal from ER to the nucleus. qPCR analysis revealed upregulation of all four genes and among these IRE1 and PERK are highly upregulated leading to 3-4 times more expression in *dcert^1^* (**Fig 5G**) whereas the levels of ATF4 transcript were not significantly changed.

### Functional conservation of human and *Drosophila* CERT

We generated transgenic flies expressing human CERT (hCERT). Western blot analysis showed that this transgene expresses a protein of ca. 75 kDa, consistent with that reported previously (**Fig 6A**). We reconstituted hCERT in the *Drosophila dcert^1^* mutant. Analysis of electrical responses to light showed that hCERT was able to rescue the reduced ERG amplitude seen in *dcert^1^* (**Fig 6B, C)**. We studied the sub-cellular distribution of hCERT using an antibody against this protein. This showed that hCERT was enriched at the base of the rhabdomere (**Fig 6D**) in a pattern consistent with the distribution of the ER-PM MCS in these cells(Yadav et al., 2015, 2018). Co-staining with the ER marker Calnexin99A showed that hCERT when expressed in adult photoreceptors was present both at the SMC as part of the ER-PM MCS and at the perinuclear ER (**Fig 6D**).

**Figure 6:**
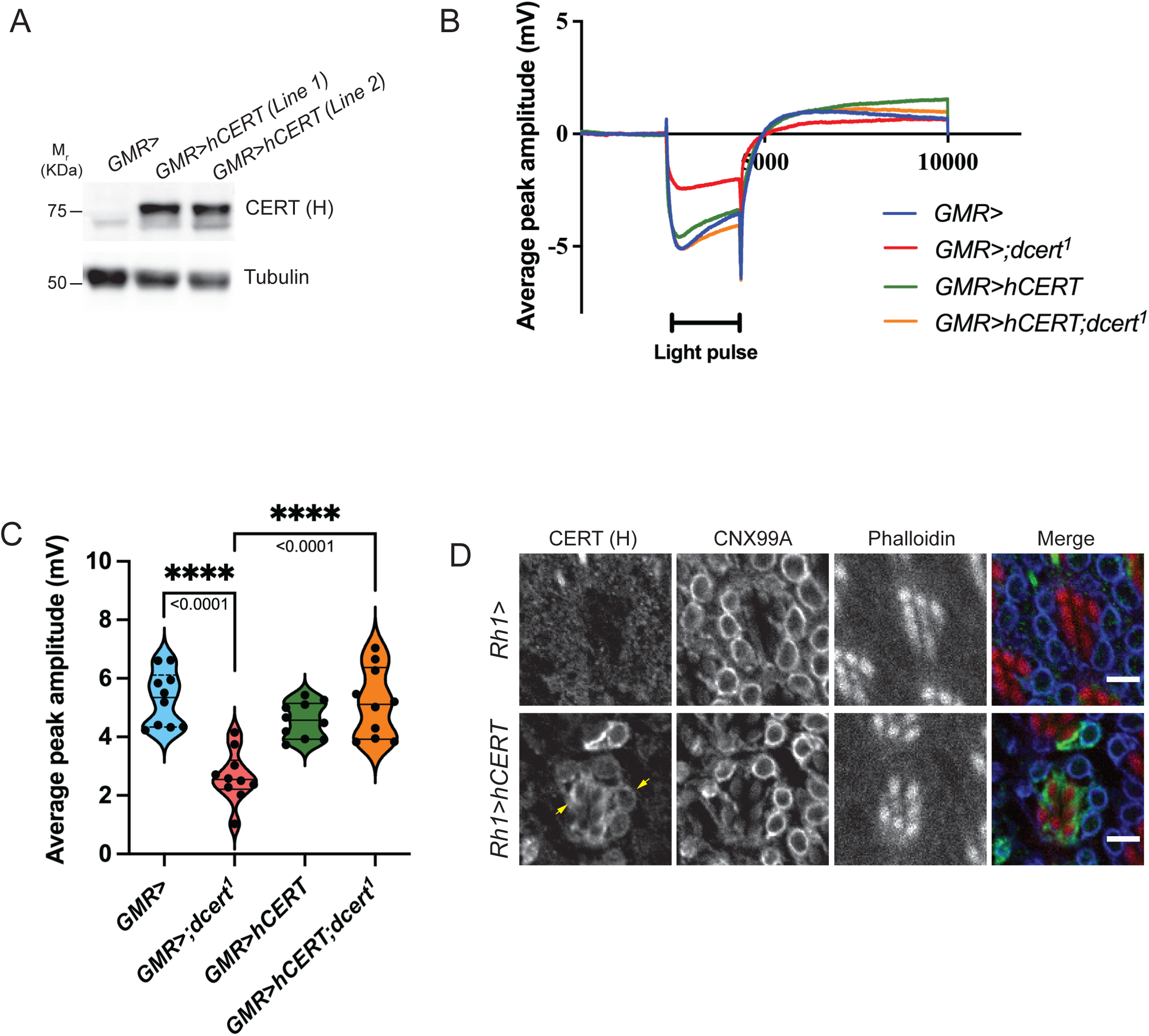
Human CERT overexpression rescues ERG defect in *dcert^1^*: (A) Western blot analysis from fly head extract of indicated genotypes is shown. The blot was probed with human anti-CERT antibody. Tubulin was used as a loading control. (B) Representative ERG traces from dark reared 0 to 1 day old flies indicating the response of control (GMR>), *GMR>;dcert^1^*, *GMR>hCERT* and *GMR>hCERT;dcert^1^* photoreceptors to 2s flash of green light. The duration of the stimulating light is shown. X axis shows the duration of recording (msec) while y axis shows the amplitude (mV). (C) Quantification of ERG amplitude of indicated genotypes. Each data point depicts a single fly tested (n=10 biological replicates). Violin plots with the mean ± SEM are shown. Statistical tests: Student’s unpaired t test. (D) Confocal images showing localization of CERT (H) in control (*Rh1>*) and *Rh1>hCERT* photoreceptors of 1-day old dark reared flies. Rh1 promoter drives the expression of hCERT only in R1-R6 photoreceptors. Yellow arrow shows the localization of human CERT at the base of the rhabdomere and at perinuclear ER colocalizing with calnexin 99A (CNX99A). Rhabdomeres are stained with phalloidin that binds to F-actin. Scale bar: 5μm.

## Discussion

Genetic analysis in multiple model systems has demonstrated the importance of CERT for physiological function. At the same time, CERT activity has been linked to the function of multiple organelles including the ER, the Golgi and PM. Given it’s role in sphingolipid biosynthesis, CERT activity is likely to affect the levels of multiple sphingolipids. To understand the mechanistic link between the biochemical function of CERT, its role in regulating organelle biology and physiological function, we studied *Drosophila* photoreceptors and found that genetic depletion of *dcert* results in a reduced electrical response to light. The ERG amplitude in *dcert* mutants progressively decreased with age, indicating a need for CERT activity in supporting lipid homeostasis at the plasma membrane and this is crucial to maintain a normal electrical response to light. Overall, these findings provide strong support for the idea that CERT activity is necessary to support normal function of G-protein coupled receptor signaling in the nervous system. Interestingly, sphingolipids are enriched in the brain (van Echten-Deckert and Herget, 2006) and have been linked to many brain diseases (Piccinini et al., 2010) although the mechanism by which they support brain structure and function is not clear. In the human nervous system, many GPCRs play functional role in diverse processes including neuromodulation through processes such as synaptic transmission and cell migration during brain development. In this context, it is interesting to note that mutations in critical residues of CERT have been noted in patients with intellectual disability. It will be interesting to explore the cellular basis of this observation and the possible link between CERT and GPCR signaling in the human nervous system. Our finding that the key phenotypes of *dcert^1^* could be rescued by hCERT position the fly photoreceptor as a model system for understanding CERT function in human brain cells and brain disorders.

Depletion of *dcert* is expected to affect the levels of multiple sphingolipids, one or more of which could contribute to the phenotypes we observed in the *Drosophila* photoreceptor system. Our biochemical analysis demonstrated elevation in the levels of several but not all ceramide species as well as a drop in the levels of PE-Cer. Although we were able to reduce the levels of ceramide by depleting SPT, the ER localized enzyme that initiates ceramide synthesis, this did not result in a complete rescue of the ERG amplitude, implying that the ER accumulated ceramide is not the sole factor underlying this phenotype. Although we noted some evidence for activation of the UPR indicating ER stress, we did not note changes in the levels or localization of the major phototransduction proteins implying that protein synthesis and trafficking is largely unaffected. It is possible that as recently reported, changes in ceramide levels may also directly induced the UPR(Kar et al., 2023; T et al., 2024). Overall, these findings may imply that elevated ceramide levels in the ER cannot solely explain the phenotypes we have observed following DCERT depletion. However, our observation that the reduced ERG amplitude could be phenocopied by depletion of CPES suggests that reduced PE-Cer may also contribute to this phenotype. Though CPES depletion also resulted in elevated ceramide levels; presumably this elevated ceramide is not accumulated in the ER since these CPES depleted flies presumably have normal CERT levels. Our analysis of *dcert^1^* photoreceptors revealed that while the levels of PM proteins such as TRP and INAD were unaffected, markers of DRMs such as contactin were reduced in *dcert^1^*. A role for sphingolipids in regulating receptor signaling at the PM has been previously proposed (Combs et al., 2013) and a previous study in *Drosophila* photoreceptors has proposed that signaling complexes at the apical PM may be organized into such domains(Sanxaridis et al., 2007). Thus, in the context of the *dcert^1^* mutant, reduced PE-Cer levels at the PM and associated reduction in DRMs likely leading to altered GPCR signal transduction and a reduced ERG response. Interestingly, a previous study in mammalian cells has shown that reduction of sphingomyelin levels without an elevation of ceramide levels alters DRMs (Fukasawa et al., 2000). Our study provides evidence of a role for CERT in this process in the context of an *in vivo* physiological process.

Due to technical difficulties, we were unable to determine the sub-cellular localization of dCERT. However, we were able to determine the sub-cellular localization of hCERT when expressed in *Drosophila* photoreceptors. Although mammalian CERT function has primarily been discussed in the context of ER-Golgi MCS, surprisingly, when expressed in fly photoreceptors, the protein was enriched to the ER-PM MCS. The ER protein VAP-A is also enriched at this location (Yadav et al., 2018) and presumably the interaction between the FFAT motif of hCERT and the MSP domain of VAP-A helps enrich it at this location. Importantly, when localized in this manner, hCERT was able to rescue the reduced ERG amplitude of *dcert^1^* implying that it is functional with respect to this phenotype at this location. Consistent with a role for CERT at this location, *dcert^1^*showed reduced ER-PM MCS and RDGB a protein selectively localized to this MCS was reduced at this location. Further, following PLC activation and depletion of PIP_2_ and PI4P at the PM, the rate of recovery in the levels of these lipids was somewhat reduced in dCERT depleted photoreceptors. These observations could also explain the reduced ERG amplitude in *dcert^1^*. However, role of dCERT in regulating phosphoinositide resynthesis is likely to be relatively modest since the reduction in ERG amplitude and rate of PIP_2_ resynthesis are much smaller compared to *rdgB* mutants(Yadav et al., 2015) and in contrast to *rdgB* mutants that undergo retinal degeneration when exposed to light(Yadav et al., 2015), *dcert^1^* photoreceptors preserve their structure on illumination (this study). In summary, this study provides compelling evidence of a role for CERT in regulating plasma membrane PE-Cer levels and signalling by GPCR.

## Acknowledgements

This work was supported by the Department of Atomic Energy, Government of India, under Project Identification No. RTI 4006 and a Wellcome-DBT India Alliance Early Career Fellowship to SM (IA/E/17/1/503653). VRN was supported by a fellowship from the Indian Council for Medical Research (2021-9971/CMB-BMS). We acknowledge the Bloomington Drosophila Stock Center (BDSC) for fly stocks. We thank the NCBS Imaging Facility, Electron microscopy and *Drosophila* facilities for support. We thank Dr. Jairaj Acharya, Dr. Susan Tsunoda, Dr. Manzoor Bhat and Dr. Armin Huber for sharing valuable reagents. We thank the NCBS Imaging, Drosophila and mass spectrometry facilities for support.

## Materials and Methods

### Fly stocks

Fly stocks were maintained in 25°C laboratory incubators with 50% relative humidity and no internal illumination. All flies were raised on standard corn meal media containing 1.5% yeast. For constant dark rearing, flies within the vials were kept in tightly closed dark boxes and maintained in the constant temperature laboratory incubator. The wild type used were *w^1118^* and Red Oregon-R (ROR). Gal4 UAS system was used for targeted expression of transgenic constructs/RNAi. The following fly alleles and insertions obtained for the experiments are described here: *dcert^1^*(Kind gift from Dr. Jairaj Acharya lab), *dCERT^BAC^*(BAC32C17,#90143 Bloomington stock), *dCERT^Deficiency^* (Df(3L)BSC732, #26830 Bloomington stock), trp>PH-PLCδ-GFP (Lab generated), UAS-P4M-GFP (Lab generated), *Lace^K05305^* (#12176, Bloomington stock), *CPES^RNAi^* (#36103, Bloomington stock), *CPES^RNAi^*(#V3594, VDRC), *SMSr^RNAi^* (#67369, Bloomington stock), *Lace^RNAi^*(#V110181, VDRC), *Lace^RNAi^* (#V21805, VDRC). All the experimental aspects of the project including animal care and use hold Institutional Biosafety Committee (IBSC) approval.

### Optical neutralization

The flies reared under experimental conditions were immobilized by cooling on ice, carefully decapitated, and fixed on the microscope slide using a drop of colorless nail varnish. The refractive index of the cornea was neutralized using a drop of immersion oil, viewed, and imaged under the 40X oil immersion objective of Olympus BX43 upright microscope (RRID:SCR_022568). The digital image acquisition and documentation were done by using CellSens software (RRID: SCR_014551).

### Electroretinogram

Anesthetized flies were introduced into truncated 200 µl disposable pipette tips such that the head protruded from the small opening. The fly’s head was then immobilized using colorless nail varnish. Two glass microelectrodes (Model no. G100F-3, Warner Instruments) filled with 0.8% NaCl solution were used for recordings such that the voltage changes were recorded by placing the experimental electrode on the surface of the eye and the reference electrode on the thorax. The flies were dark adapted for 5 min prior to ERG recordings followed by 2-s flash of green light stimulus, with 10 stimuli (flashes) per recording and 15 s of recovery time between two subsequent flashes. Green light stimulus was emitted using an LED light source to within 5 mm of fly’s eye through a fiber optic guide. Voltage changes were recorded using pCLAMP 10.7 and amplified using DAM50 amplifier (SYS-DAM50, WPI, Florida, USA). Data analysis was done using Clampfit 10.7 (Molecular Devices, California, USA). Graphs were plotted using GraphPad Prism software (RRID: SCR_002798).

### Western blot

Age-matched flies were decapitated, and the heads were homogenized in 2× Laemmli sample buffer followed by boiling at 95°C for 5 min. For rhodopsin blot, the protein samples were processed differently. The fly heads were snap-frozen in liquid nitrogen and stored at 80°C for a period of 2 days. These head samples were homogenized in 2× Laemmli sample buffer followed by boiling at 37°C for 30 min. Protein extracts from fly heads were separated using SDS–PAGE and transferred onto nitrocellulose filter membrane [Hybond-C Extra; (GE Healthcare, Buckinghamshire, UK)] using wet transfer apparatus (Bio-Rad, California, USA). The membrane was blocked using 5% Blotto (sc-2325, Santa Cruz Biotechnology, Texas, USA) in phosphate buffer saline (PBS) with 0.1% Tween 20 (Sigma-Aldrich) (PBST) for 2 h at room temperature (RT). Primary antibody incubation was done overnight at 4°C using appropriate antibody dilutions: chicken anti-dCERT (kind gift from Dr. Jairaj Acharya lab), 1:1000; rat anti-RDGB (lab generated), 1:4,000; mouse anti-rhodopsin (DSHB Cat# 4c5, RRID:AB_528451), 1:250; rabbit anti-Arrestin2 (kind gift from Dr. Armin Huber lab), 1:1,000; rat anti-TRP (lab generated), 1:4,000; rabbit anti-norpA (kind gift from Dr. Armin Huber lab), 1:1,000; rabbit anti-INAD (kind gift from Dr. Susan Tsunoda lab), 1:1,000; rabbit anti-TRPL (kind gift from Dr. Armin Huber lab), 1:1000; anti-dPIP5K (lab generated), 1:1000; anti CERT (human, PA5-113546, Invitrogen), 1:1000 and mouse anti-tubulin-E7c (DSHB Cat# E7, RRID:AB_528499), 1:4,000. Following this, the membrane was washed in PBST (3 times for 10 mins each) and incubated with 1:10,000 dilutions of appropriate secondary antibody (Jackson ImmunoResearch Laboratories, Pennsylvania, USA) coupled to horseradish peroxidase at RT for 2 h. Three PBST washes (10 mins each) were given followed with one 10 min 1XPBS wash, and the blots were developed with ECL (GE Healthcare) and imaged using LAS 4000 imageQuant (RRID: SCR_014246) (GE Healthcare).

### Fluorescent pseudopupil

To monitor changes in PIP_2_ levels at the microvillar PM in live flies, the PIP2 biosensor PH-PLCd coupled to GFP driven by the transient receptor (*trp)* promoter of flies was used (Yadav et al, 2015). The flies were made insentient and were immobilized using the same protocol as in ERG recordings. The pseudopupil formed from the summed fluorescence of approximately 20–40 adjacent ommatidia were focused and imaged using the 10× objective of Olympus IX71 microscope (RRID:SCR_022185). The program created using the software Micromanager captured time-lapse images of the pseudopupil by collecting fluorescence emitted from the eye when GFP was stimulated by a 90-ms flash of blue light. Prior to the recordings, the flies were dark adapted (resting conditions) for 6 min during which the probe binds to PIP_2_ and localizes to the microvillar PM. Following a flash of blue light, the PLC8 activity triggers the hydrolysis of PIP_2_, and thereby, the probe is displaced from the PM and this leads to the loss of the pseudopupil. The central UV-sensitive photoreceptor is unresponsive to blue light and thereby retains the probe at the apical PM. Subsequent red light illuminations following the blue light stimulus after each time-lapse image acquisition hastens the retrieval of the probe back to the PM. Mean fluorescence intensity indicating the basal PIP_2_ pools and the PIP_2_ recovery kinetics were calculated using ImageJ from NIH (Bethesda, MD, USA) (RRID:SCR_003070). Quantification of DPP fluorescence intensity was done by measuring the intensity values per area of the pseudopupil. Similarly, flies expressing the P4M-GFP probe (PI4P biosensor) were subjected to the same protocol for monitoring PI4P levels. P4M-GFP expression was driven by an eye-specific promoter GMR using Gal4-UAS system.

### RNA extraction and qPCR

RNA isolation was done from Drosophila heads using TRIzol reagent (Invitrogen) followed by treatment with amplification grade DNAse I (Invitrogen). cDNA synthesis was done using the Superscript II RNAse H Reverse Transcriptase (Invitrogen) and random hexamers (Applied biosystems). Mentioned primers were designed at the exon– exon junction of the genomic region satisfying the parameters recommended for qPCR primer designing, and the run was performed in the Applied Biosystem7500 Fast Real Time PCR instrument using Ribosomal Protein 49 (RP49) primers as the control primers flanking the housekeeping gene region. Triplicates of each sample were measured to ensure the consistency of the data. The following primers were used for qPCR:

RP49 forward: CGGATCGATATGCTAAGCTGT

RP49 reverse: GCGCTTGTTCGATCCGTA

PI4KIIIα forward: GAGGAACAGATCACGGAATGGCG

PI4KIIIα reverse: CCTCCTCATCAATGATCTCCGCG

PERK forward: GAGGCTCGAACTTTGGC

PERK reverse: CTTGCGATCCTCTTCTTCC IRE1

forward: CCATCGATCCGGTGACC IRE1

reverse: GGATCGGGCAGAAAGTG HSP7

forward: CGTATTCCTGCGTTGGTG

HSP70 reverse: GTGGTCAACTGATTCTTGGC ATF4

forward: GCCAAAACCCGTGCTC

ATF4 reverse: CTCGGGCAATTTCGCTG

### Immunohistochemistry

Heads were carefully removed, placed on double sided tape and immersed with fixing solution (4% paraformaldehyde in PBS with 1 mg/ml saponin) at room temperature for 20 mins. Heads were washed two times with ice cold 1x PBS and retinae were dissected in 1x PBS. Retinae were fixed again on ice for 20 mins. Post fixation, the samples were washed in PBS with 0.3% Triton X-100 (0.3% PBTX) followed by incubation with blocking solution (5% fetal bovine solution in PBTX) for 2 h at RT. The samples were then incubated overnight with the respective antibody [anti-RDGB (lab generated), 1:300; anti-TRP (lab generated), 1:200; anti-INAD (kind gift from Dr. Susan Tsunoda), 1:200; anti-Rh1 (DSHB, 4C5c), 1:25 and anti CERT (human, PA5-113546, Invitrogen), 1:100] at 4°C. After this, samples were washed thrice with 0.3% PBTX and incubated with appropriate secondary antibody [Alexa Fluor 488 anti-rabbit (Thermo Fisher Scientific Cat# A-11034, RRID:AB_2576217), Alexa Fluor 488 anti-rat (Thermo Fisher Scientific Cat# A-11006, RRID:AB_2534074); 1:300 dilution] for 4 hrs at RT. Along with the secondary antibody incubation, Alexa Fluor 633–Phalloidin (Invitrogen, A22284; 1:2000) was used to mark F-actin. Samples were washed thrice with 0.3% PBTX, followed by one final wash in PBS and were mounted with 70% glycerol in 1× PBS. The whole-mounted preparations were imaged under 60 × 1.4 NA objective, Olympus Confocal Laser Scanning Microscope Fluoview FV3000 (RRID:SCR_017015).

### Quantification of RDGB intensity

Using the freehand line on the tool bar in ImageJ (RRID:SCR_003070) the line was drawn across the RDGB fluorescence at the base of the rhabdomere. Using plot profile tool under Analyze pixel intensity graph was generated. By selecting list in the plot profile highest gray value was recorded. Intensity profile from 30 photoreceptors (5 ommatidia) from single retina forms one biological replicate. In each experiment at least 3 biological replicates with total 90 photoreceptors (from 15 ommatidia) corresponding three independent retinae were taken for quantification. Graphs were plotted using GraphPad Prism software (RRID: SCR_002798).

### Electron microscopy

The fly heads of mentioned genotypes were cut sagittal and immersed in 2% osmium tetroxide, kept at 4°C for 1 h followed by incubation at 40°C for 4 days (Heads will remain afloat). Heads were washed with distilled water, stained enbloc with uranyl acetate (0.5% in distilled water) for 3 h. After washing with distilled water, heads were subjected to dehydration step gradually starting with 50% ethanol molecular grade (2 times, 10 mins each), 70% ethanol (3 times, 10 mins each) and 100% ethanol (3 times, 10 mins each). After the ethanol wash, all heads will sink to the bottom of Eppendorf tube. After ethanol wash heads were treated with ethanol: propylene oxide (1:1 by volume) for 15 mins on rotor followed with only propylene oxide (5 mins). Afterwards heads were immersed in propylene oxide : epoxy resin (3:1 by volume) for 30 mins on rotor followed by propylene oxide: epoxy resin (1:1 by volume) for 30 mins on rotor followed by propylene oxide: epoxy resin (1:3 by volume) for 30 mins on rotor and finally only in epoxy resin overnight. Next day heads were embedded in resin. Ultrathin sections of 60 nm were cut, and grids were subjected to poststaining with 2% uranyl acetate (in 70% ethanol) and Reynold’s lead solution. Sections were imaged at 120 KV on a Tecnai G2 Spirit Bio-TWIN (FEI) electron microscope (RRID:SCR_022981).

### Scoring MCS density

For scoring MCS density/photoreceptor cell, a total of 30 cells were taken to conduct analysis for R1-R6 photoreceptors. Using free hand line tool of ImageJ (RRID:SCR_003070), length of MCS (μm)/the total length of the base of the rhabdomere (μm) was calculated. Fractions of MCS coverage were multiplied with 100 to show the percentage. Groups were compared using the GraphPad Prism software (RRID: SCR_002798).

### Separation of detergent resistant membrane (DRM)

DRMs were separated as described in Kunduri et al., PNAS 2019 with few modifications. 400-500 heads from day 1 old flies were snap frozen in liquid nitrogen and separated. Heads were then collected in the homogenizer (Wheaton, 7ml) and homogenized first with loose pestle followed with tight pestle in 2 ml of 5 mM EDTA,10 mM Tris pH 7.5, 0.25 M sucrose with protease inhibitor cocktail (Roche). Later the volume was adjusted to 4.5 ml and homogenized. To the final homogenate 0.5 ml of 1M KCL was added to make final concentration of 100mM. The homogenate was centrifuged at 3000 rpm at 4°C for 10 min on tabletop centrifuge. The resulting supernatant was centrifuged at 100,000 g for 1 hour at 4°C (Beckman Coulter Optima Max-XP Tabletop Ultracentrifuge (RRID:SCR_025704), TLA 100.3 rotor). Resulting pellet was resuspended in 200 μl of buffer (10 mM Tris pH 7.5, 5 mM EDTA, 0.25 M sucrose, 100 mM KCl and protease inhibitor cocktail) and volume was made upto 1 ml. 50 μl was taken for protein estimation using Bradford (Sigma) and input protein sample. The resuspended membrane (950 μl) was mixed with 950 μl of buffer having 2% Triton X100 and protease inhibitor and incubated on ice for 30 min. the solubilised membranes were mixed with 2 ml of 80% sucrose in buffer and transferred to ultra-clear centrifuge tubes (14x 95 Ultra Clear Beckman) and layered with 4 ml of 30 % and 4 ml of 5 % sucrose to form a gradient. The gradient tubes were centrifuged at 39,600 rpm for 16 h at 4°C (Optima L-100K Beckman Coulter, SW40Ti rotor). Following centrifugation, 1 ml fractions were collected from top to bottom.

### Western blotting

20 μl protein samples were mixed with 10 μl of 2 X Lamelli buffer and incubated at 95°C for 5 minutes except for Rh1 which was incubated at 37 °C for one hour. Equal amount of protein sample was loaded and separated on SDS-PAGE, transferred to nitrocellulose membrane using wet transfer system for 2 hours at 100V. The membrane was blocked using 5% Blotto (sc-2325, Santa Cruz Biotechnology, Texas, USA) in phosphate buffer saline (PBS) with 0.1% Tween 20 (Sigma-Aldrich) (PBST) for 2 h at room temperature (RT). Primary antibody incubation was done overnight at 4°C using appropriate antibody dilutions: guinea pig anti contactin (Kind gift from Dr. Manzoor Bhat lab), 1:2000; mouse monoclonal Na+/K+-ATPase antibody (DSHB Cat# a5, RRID:AB_2166869), 1:500.

Following this, the membrane was washed in PBST (3 times for 10 mins each) and incubated with 1:10,000 dilutions of appropriate secondary antibody (Jackson ImmunoResearch Laboratories, Pennsylvania, USA) coupled to horseradish peroxidase at RT for 2 h. Three PBST washes (10 mins each) were given followed with one 10 min 1XPBS wash, and the blots were developed with ECL (GE Healthcare) and imaged using LAS 4000 imageQuant (RRID: SCR_014246) (GE Healthcare).

### Sphingolipid isolation

Flies were flash frozen in liquid nitrogen and heads (150 per biological replicate) or retinae (300 per biological replicate) were collected in Precellys vials with beads. Sphingolipids were isolated with modification in previously described (1,2). Drosophila head and retina samples were homogenized by probe sonicator (Sonics VibraCell VCX130, USA) in LC-MS grade water. An aliquot was taken for protein estimation, which was done by Bicinchoninic Acid (BCA) kit. Suspension along with Internal standard (Cer/Sph mixture II) was added to 1.5 ml of chloroform: methanol (1:2 v/v) in fresh borosilicate tube and incubated overnight. 1M KOH was added, briefly sonicated, and further incubated at 37 degrees shaking incubator for 2 hours to cleave interfering glycerolipids. Samples were neutralized with glacial acetic acid, centrifuged to remove debris and supernatant transferred to other tube. Insoluble debris were further re-extracted with 0.5 mL of chloroform: methanol (2:1 v/v) and supernatant was transferred to previous tube. It was sequentially mixed with 1 mL chloroform and 3.5 mL water and centrifuged. After centrifugation lower layer was transferred, upper phase was re-extracted and subsequently lower phase was dried under nitrogen gas at 48 degrees.

### Liquid chromatography-Mass spectrometry

Lipids were analyzed (Goh and Guan, 2021) by multiple reaction monitoring (MRM) approach using high pressure UHPLC liquid chromatography (ExionLC AC, SCIEX, USA) coupled to a hybrid triple quadrupole/linear ion trap mass spectrometer (4500 QTRAP, SCIEX, USA). Components were separated by Acquity UPLC Beh C18, 2.1 x 50 mm column (Waters®) with a particle size of 1.7 μm using solvent A (water: Formic acid (99.8:0.2 v/v) with 5 mM ammonium formate) and solvent B (Methanol: Formic acid (99.8:0.2 v/v) with 5 mM ammonium formate). Dried samples were reconstituted in 300 uL buffer A: B (2:8 v/v), centrifuged at 10000 rpm for 10 min and supernatant was transferred to autosampler vials for further analysis. For separation solvent A and solvent B is maintained at 20:80 ratio for 0-6 min, followed only solvent B (100%) for 6-17.5 min, and finally a wash with mixture of solvent A and B in 20:80 ratio from 18.1 to 20 min before the next run.

### Data analyses

Sphingolipids were monitored using MRM mode. Acquisition was done using Analyst 1.6.3 software and peak area for all analytes was estimated into MultiQuant 3.0.2 Multiquant 3.0.2 software (AB SCIEX, USA) for data analysis to quantitate lipid species. A standard curve was generated for quantitation (pmol/mg) of sphingolipids. To quantify analyte in each sample, the area under the peaks for both analyte and spiked internal standard was estimated, and ratio of each analyte with internal standard was determined. These ratio values were used to quantitate the lipid species. Students t-test was used to check the statistical significance between different groups.

