## Supplementary figures and images for "Ceramide transfer protein regulates G-protein coupled phospholipase signalling in *Drosophila* photoreceptors"

### supplementary figures

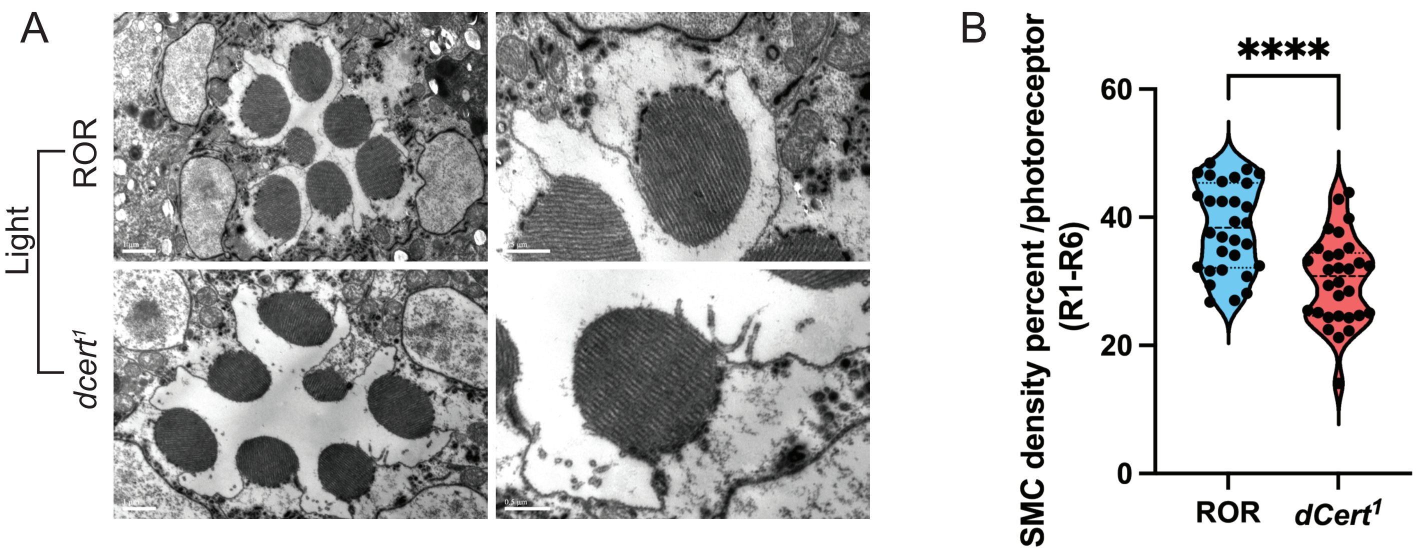


**Supplementary Figure 1**

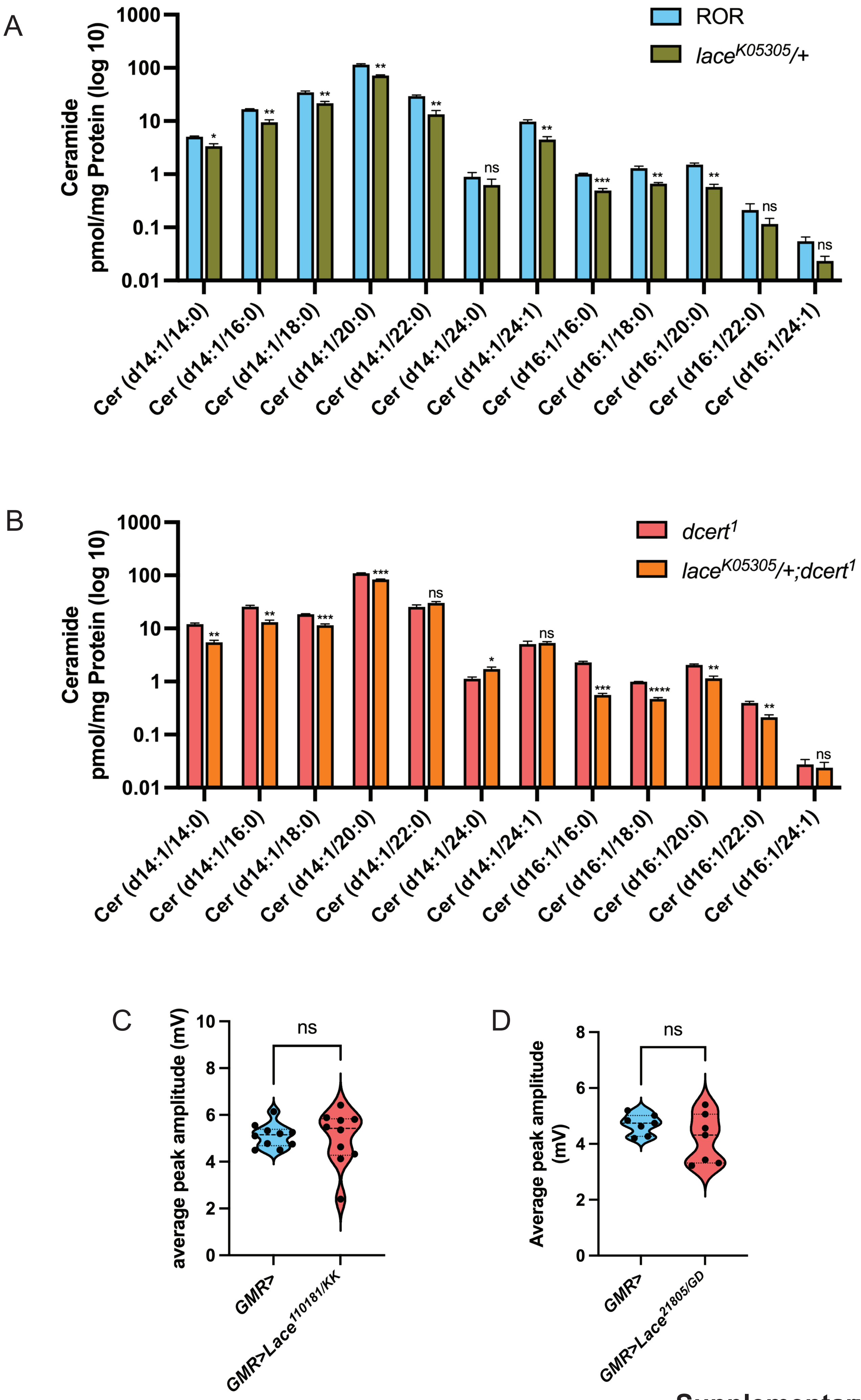


**Supplementary Figure 2**

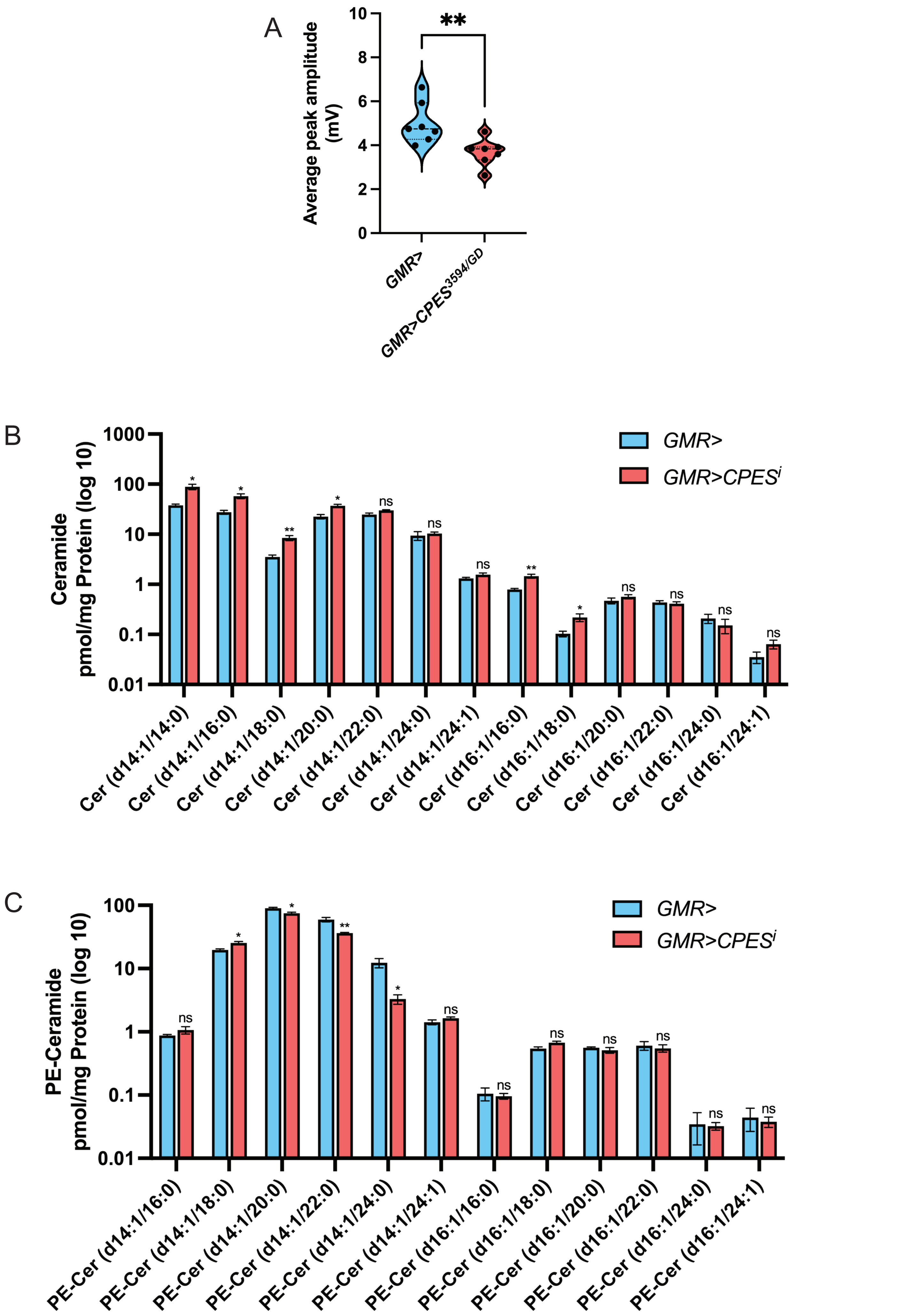


**Supplementary Figure 3**
